# Spectral decomposition of local field potentials uncovers frequency-tuned gain modulation of working memory in primate visual system

**DOI:** 10.1101/2025.07.28.667125

**Authors:** Majid Roshanaei, Mohammad Reza Daliri, Zahra Bahmani, Kelsey Clark, Behrad Noudoost

## Abstract

Working memory has been shown to modulate visual processing in a variety of ways, including changes in visual response gain, oscillatory power, spike timing, and phase coding of information^1–3^. Here we probe working memory’s influence on various oscillatory components within visual areas, using the Maximal Overlap Discrete Wavelet Transform (MODWT) technique to decompose the local field potential(LFP)^4^; this method allows single-trial quantification of the properties of various oscillatory components. We examine the impact of spatial working memory on visual processing within the extrastriate middle temporal (MT) visual area of rhesus macaque monkeys, comparing the responses in MT when remembering a location inside or outside the receptive fields of the neurons being recorded. Using traditional bandpass filtering, we replicate previous reports that working memory enhances visual responses, low-frequency oscillatory power, spike-phase locking, and phase coding of visual information. Applying the MODWT method, we find that working memory modulates both oscillatory power and the precise frequency of oscillations within a general frequency band. The precise frequency of several lower frequency components (alpha, theta, and beta frequencies, but not gamma or high gamma) correlates with visually-evoked firing rates of MT neurons on a trial-by-trial basis. This relationship between firing rate and oscillatory component frequency is maintained across memory conditions, indicating a close link between the neural mechanisms driving oscillatory frequency and firing rates.

## Introduction

Working memory (WM) is a fundamental cognitive process that enables the temporary storage and manipulation of information to guide goal-directed behavior^5,6^. While WM has been extensively studied in the context of prefrontal and parietal cortices, it also influences sensory processing^7,8^. WM can bias neural activity in primary visual cortex (V1) through feedback signals from higher-order areas, enhancing the representation of task-relevant stimuli^9^. WM can modulate sensory representations in extrastriate visual cortices, such as the middle temporal area (MT), enhancing the gain of visual signals^2,10^. WM also changes oscillatory activity in these areas: in MT, WM alters oscillatory power and spike-phase locking (SPL)^2^, and in V4 there is a WM-driven change in oscillatory power, SPL, and phase coding which depends on prefrontal activity ^3^.

Many studies (including our own prior work^2,3^) use a bandpass filter to isolate specific frequency bands from the local field potential (LFP), then look for changes in power or spike-phase locking in each of these bands, averaged across trials within a condition. While this approach has revealed many changes in oscillatory measures due to cognitive factors such as attention^11–14^ and working memory^2,3,9,15^, this method has some limitations, including the need for experimenter-selected frequency bands and the loss of information about the variability of oscillations across or within trials.

Here we use the Maximal Overlap Discrete Wavelet Transform (MODWT) technique to decompose the LFP^16^. The advantages of MODWT, compared to simple bandpass filtering, are that it 1) separates aperiodic and oscillatory components, 2) does not require a priori specification of frequency bands, and 3) operates at the level of individual trials. Several studies have employed MODWT for analyzing LFPs, particularly due to its advantages in handling non-stationary neural signals. Savolainen demonstrated that inter-frequency power correlations in LFPs, quantified through wavelet transforms, reflect functionally significant interactions between neural ensembles, with distinct coupling patterns emerging during different cognitive states^17^. This approach proved particularly valuable because wavelet methods preserve the temporal precision needed to detect brief but behaviorally relevant coupling events. Building on this, Pinzuti et al. developed a wavelet-based framework for spectrally-resolved information transfer measurement, showing that information flows preferentially between specific frequency bands during sensory processing^18^. Their method overcame limitations of traditional coherence analyses by using the wavelet transform’s adaptability to non-stationary signals. Most recently, they applied information-theoretic measures to wavelet-decomposed LFPs, revealing layer- and frequency-specific changes in cortical information processing under anesthesia^20^. Their findings highlight how wavelet methods can disentangle simultaneous changes across different cortical layers and frequency bands - a crucial advantage over conventional spectral analysis. Together, these studies establish wavelet transforms (particularly MODWT due to its shift-invariance) as essential tools for investigating the multi-scale, dynamic nature of neural information processing. We examine how a top-down WM signal alters these oscillatory components within extrastriate visual cortex, and the relationship between these oscillations and firing rate responses of neurons.

To test the influence of WM signals on visual processing, we recorded LFP and spiking activity from MT during a spatial working memory task with visual probes. Consistent with previous reports in MT and V4^1–3^, with traditional bandpass filtering of LFPs we find that WM modulates the magnitude of visually-evoked responses, oscillatory power, spike timing relative to these oscillations, and phase coding in MT. Applying the MODWT method to this data, we see that WM also alters the exact frequency of oscillations within one band, with oscillation frequency for several components shifting slightly higher at sites corresponding to the location held in WM. Across trials, this frequency is correlated with the firing rate response to visual stimuli. Linear regression shows that the relationship between component frequency and firing rate is similar regardless of the memory condition. This correlation indicates a tight link between the neural circuit mechanisms driving oscillatory frequency and firing rate modulations during WM.

## Results

This study investigates the modulation of oscillations and visual responses by working memory in the extrastriate cortex. LFP oscillations and spiking activity in area MT of two monkeys were analyzed during a memory-guided saccade task with visual probes. Recordings were made using 16-channel linear array electrodes, resulting in data from 131 isolated single units and 256 LFP channels across 16 recording sessions. As illustrated in Figure 1A, the experimental task involved presenting a monkey with a visual cue, which it had to remember throughout a delay period, then making a saccadic eye movement to the memorized location to receive a reward. During the fixation and delay periods, 200-ms visual probes were presented near the neuron’s receptive field. Each neuron’s receptive field was determined based on its firing rate response to a 7×7 matrix of probes presented during the fixation period (see Methods). The memorized location could be in various positions relative to the neuron’s receptive field, with three possible cue locations in the same hemifield as the recorded neuron (the nearest to the center of the receptive field of the neuron is referred to as the IN condition), and one cue located in the opposite hemifield (referred to as the OUT condition). The receptive field of an example neuron is shown in figure 1B. There was significant power modulation for the memory period (2400-3000ms) compared to the pre-stimulus fixation period in the αβ band, as demonstrated by the scatter plot in Figure 1D (power change_IN_= 74.07 ± 0.99%, power change_OUT_= 71.27 ± 0.94%, *p* < 10^-3^). Furthermore, significant power modulation was also observed in the theta band (%power change_IN_= 110.64 ± 1.45, %power change_OUT_= 100.15 ± 1.75, *p* < 10^-3^; see Figure S1A), while no significant effect was detected in the gamma band (%power change_IN_= 37.40 ± 0.54, %power change_OUT_= 36.84 ± 0.66, *p* = 0.623; see Figure S1B). Given recent reports of αβ phase coding in V4 during WM [ref Parto 2024], we looked for changes in the phase coding of probe location between IN and OUT conditions: 131 MT neurons were examined for mutual information between 49 probe stimuli and αβ oscillation phases at spike times (see method). As shown in Figure 1E, a significant increase in phase-coded mutual information for probe location was observed during the memory IN condition compared to the OUT condition (MI_IN_ = 0.062±0.007, MI_OUT_ = 0.019±0.007, p < 10^-3^). This mirrors the results reported in V4, and indicates that working memory enhances the information about incoming visual stimuli conveyed via αβ phase at spike times across multiple visual areas. In addition to examining phase code modulation, the firing rate differences between memory IN and OUT conditions were analyzed to assess gain modulation. As shown in Figure 1F, the average probe-evoked response during memory IN trials was slightly but significantly different from the OUT condition for 131 MT neurons (firing rate_IN_ = 7.61±0.64, firing rate_OUT_ = 7.29±0.58, p = 0.038). These results replicate the WM-driven changes in visual responses and oscillatory power reported in MT^1,2^, and expand previous findings of WM-driven phase coding in V4 to MT^3^.

**Figure 1.**
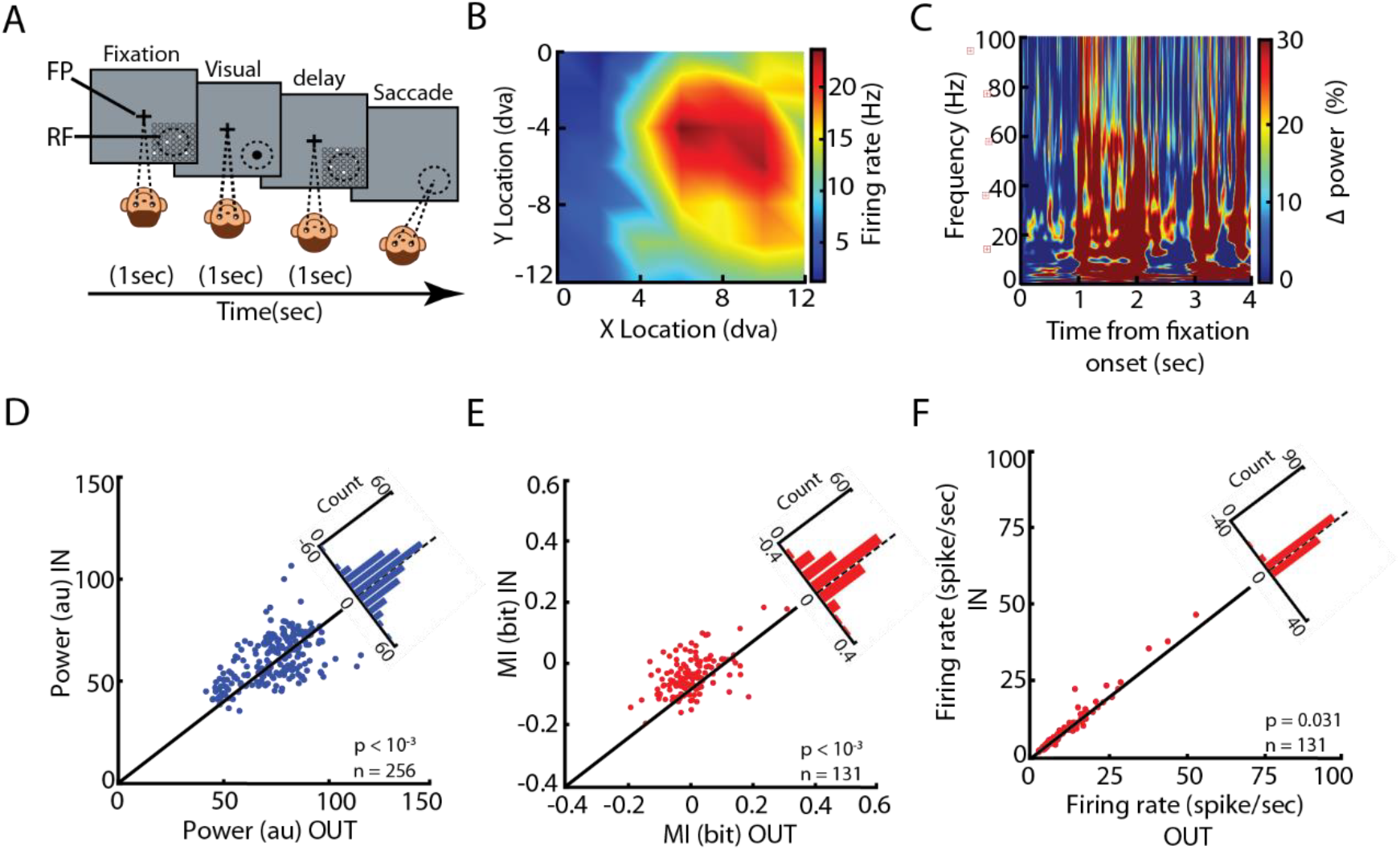
WM task, power spectrum, phase code and gain modulation. (A) The animal performed an MGS task in which visual probes appeared during the 1-s fixation period and the 1-s delay period. The probe (white circle) was a brief (200 ms) small (∼1 dva) visual stimulus presented in a 7 × 7 grid of possible locations (open white circles, shown here for illustration only and not present on the screen). In each trial, four probes were presented in succession during both the fixation and memory periods, with an inter-probe interval of 200 ms. This 7 × 7 grid of probes was positioned to overlap with the RF of the recorded neuron based on the preliminary RF mapping. The location of the remembered target could vary with respect to the RF of recorded neurons. (B) The receptive field of a sample MT neuron. Plot shows firing rate in response to probes appearing at various location. (C) Power change (relative to pre-stimulus baseline) over time and frequencies for the sample recording channel as the neuron shown in B. (D) Scatter plot of mean power change in αβ band across 256 recorded channels for memory IN and OUT condition. The diagonal histogram shows the distribution of differences in power. (E) Across 131 MT neurons, the mutual information between the 49 different probe stimuli and the phases of ongoing αβ oscillation at the time of spikes showed a significant increase during memory IN compared with memory OUT. The scatterplot shows the MI for each MT neuron during memory OUT (x axis) and during memory IN (y axis). The diagonal histogram shows the distribution of differences in MI. (F) Visually evoked firing rates increased during the memory IN period. The scatterplot shows visually evoked spiking activity for the optimal probe of each neuron in the memory IN versus memory OUT. The diagonal histogram shows the distribution of changes in the visually evoked spiking activity.

Next, the LFP signals were decomposed using the MODWT method. A key requirement of the MODWT process is the specification of the number of decomposition levels, which significantly influences the resolution and interpretability of the resulting components. To determine the appropriate number of decomposition levels, a method described by Donoghue et al. (2020)^4^ was utilized. This method involves analyzing the power spectral density (PSD) of the single trial LFP signal by fitting a model that separates the periodic (oscillatory) and aperiodic (non-oscillatory) components. In Figure 2A, the original PSD of a single LFP trial is represented by the solid black line, while the solid red line and dashed blue line represent the periodic and aperiodic components, respectively (R^2^=0.989). This analysis allows for the identification of key spectral features, including the number and specific frequencies of peaks in the PSD. The analysis was conducted on trials from 16 sessions, each of which contained recordings from 16 channels. As shown in the top panel of Figure 2B, the distribution of the number of peak frequencies across trials within these sessions indicates that most trials exhibit six distinct peak frequencies. However, one of these frequencies corresponds to the DC component (0 Hz), which is treated as the approximation component in the MODWT framework. Based on this observation, five levels of decomposition were selected for further analysis, excluding the DC component to focus on meaningful frequency bands. This approach ensures that the number of decomposition levels corresponds to the spectral properties of the data, enhancing the accuracy and relevance of subsequent analyses. Additionally, the bottom panel of Figure 2B illustrates the distribution of frequencies across these sessions. The schematic representation of the MODWT method is presented in Figure 2C, illustrating the five levels of decomposition applied to the LFP signal (see method). In this process, the original signal is iteratively decomposed into approximation and detail coefficients at each level. The approximation coefficients capture the lower-frequency components of the signal, while the detail coefficients represent the higher-frequency oscillatory features. The decomposition proceeds through five levels, with each level focusing on progressively finer frequency bands. The final approximation coefficients at level five represent the lowest-frequency components, including the DC component, which corresponds to the overall trend or baseline of the signal. Meanwhile, the detail coefficients across the five levels collectively capture distinct higher-frequency bands, enabling the identification of oscillatory features within the signal. Figure 2C provides a visual summary of this process, highlighting how the MODWT method systematically separates the signal into its frequency components over the five decomposition levels. This hierarchical representation enhances the ability to analyze spectral and temporal characteristics of the LFP data in a structured manner. The MODWT method was applied to decompose sample trials (0 to 3000 ms) into five distinct detail components and one approximation component, as shown in Figure 2D. The first subplot displays the original signal (red line), while the subsequent subplots illustrate the decomposed detail components and the approximation component. The peak frequency associated with each component is indicated in the title of its respective subplot. After applying MODWT to all trials across all channels, the peak frequency of each decomposed component was extracted for every trial. Figure 2E provides a comparative visualization of the peak frequency distributions for each component across 256 channels under two experimental conditions: IN and OUT. The figure highlights distinct differences in the frequency characteristics between the two conditions. Figure 2E consists of six subplots, each representing a scatter plot for one component. In each subplot, the peak frequencies for the 256 channels are compared between the IN condition (y-axis) and the OUT condition (x-axis). The scatter plots reveal distinct patterns: higher peak frequencies are observed for the IN condition in the alpha, beta, low gamma and delta bands, whereas the OUT condition exhibits higher peak frequencies in the high gamma band (frequency_delta-IN_=1.41±0.01, frequency_delta-OUT_=1.40±0.01, p=0.012; frequency_theta-IN_=5.96±0.01, frequency_theta-OUT_=5.97±0.0q, p=0.061; frequency_alpha-IN_=11.25±0.04, frequency_alpha-OUT_=11.03±0.03, p<10^-3^; frequency_beta-IN_=20.49±0.04, frequency_beta-OUT_=20.29±0.04, p<10^-3^; frequency_low gamma-IN_=39.70±0.06, frequency_low gamma-OUT_=39.41±0.06, p<10^-3^; frequency_high gamma-IN_=79.59±0.10, frequency_high gamma-OUT_=80.17±0.12, p<10^-3^). These results suggest differential modulation of neural oscillations by working memory, with specific frequency bands being selectively enhanced during the IN condition.

**Figure 2.**
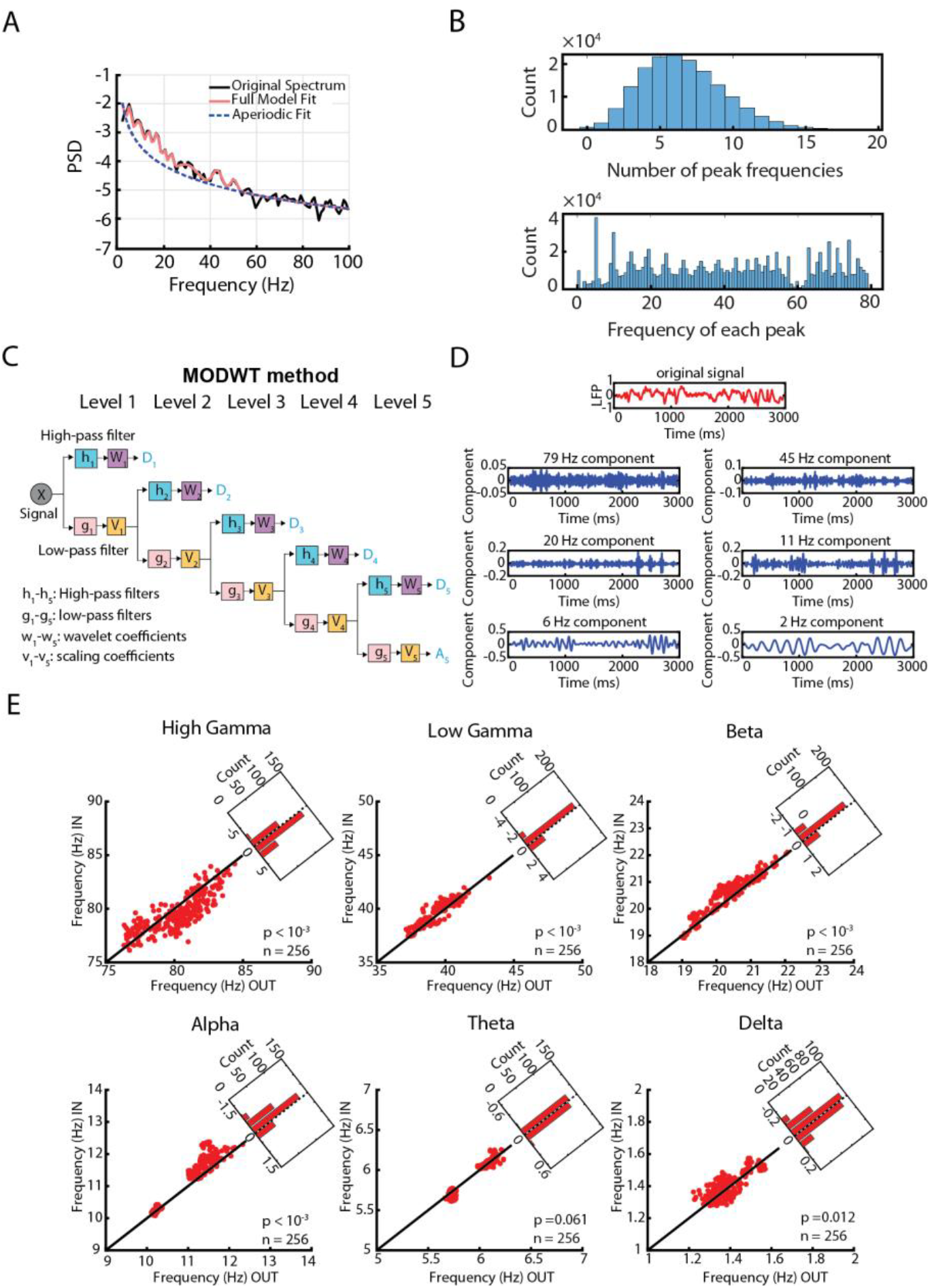
Analysis of Peak Frequencies, Decomposition Method, and Component Frequencies. (A) The original power spectral density of a single trial (black solid line), fitted power spectral density (red solid line), and aperiodic component of the power spectral density (dashed blue line). (B) The top panel shows a histogram of the number of peak frequencies in trials across all recording sessions, while the bottom panel displays the detected frequencies. (C) Schematic Diagram of a Five-Level MODWT Method. This schematic illustrates the hierarchical decomposition process of a time series signal using MODWT, depicting five successive levels of transformation. Each level shows the corresponding wavelet and scaling coefficients, highlighting how the signal is progressively decomposed into finer frequency components and its approximation. (D) Decomposition of a single trial using MODWT. The first subplot (red line) shows the original signal, and the remaining subplots display the decomposed components, with the peak frequency of each component indicated in the title of each subplot. (E) Scatter plot of the frequency of each of the six components for memory IN and OUT conditions across 256 recording channels.

After the decomposition process, power differences for each frequency component during the memory period (2400–3000 ms) between IN and OUT conditions were analyzed for the sample neuron depicted in Figure 1B, as shown in Figure 3A. This analysis revealed prominent modulations in the power of the alpha, beta and theta components, aligning with previously discussed findings. A cross-sectional view of power differences for each component (see Methods), illustrated in Figure 3B, reveals sustained activity in the alpha, beta, and theta components during the memory period, highlighting distinct patterns of power modulation across these frequency bands. The power change for the theta component showed a significant deviation from baseline (0-50 ms) between 2572 ms and 2858 ms (time bin-sliding 50ms bin for comparison with Wilcoxon signed rank). Similarly, significant changes were observed for the beta component from 2460 ms to 3000 ms and for the alpha component from 2432 ms to 3000 ms. To investigate the influence of frequency on visual signal gain, visual probes were ranked from 1 to 49 based on their evoked responses during the memory IN period. The firing rates associated with these ranked probes were categorized into three frequency groups (high, middle, and low) for each frequency band, based on the frequency of the decomposed components of the corresponding LFP signal, as illustrated in Figure 3C (see Methods). This result, derived from the same sample neuron shown in Figure 1B, demonstrates that both the frequency of the components and the optimality of the probes significantly impact signal gain. The analysis indicates that firing rates are significantly modulated by frequency in the theta, alpha, and beta components (p<10^-3^, f-statistic_theta_= 32.97, f-statistic_alpha_= 61.78 and f-statistic_beta_ = 34.43, ANOVA), while no significant modulation was observed in the delta, low gamma, and high gamma components (p>0.05, f-statistic_delta_= 1.22, f-statistic_low gamma_= 5.94 and f-statistic_high gamma_ = 4.29, ANOVA). Additionally, firing rates were significantly modulated by probe optimality only in the low gamma component (p=0.021, f-statistic_low gamma_= 2.61, ANOVA). To quantify these relationships, a linear regression model was applied to the frequency-divided firing rate data, as illustrated in Figure 3D (mean R^2^ = 0.64 ± 0.05). The fitted model demonstrates that frequency significantly contributes to the modulation of neuronal gain in the theta, alpha, and beta components (p<10^-3^, t-statistic_theta_=6.11, t-statistic_alpha_=10.19 and t-statistic_beta_=8.30). In contrast, probe optimality significantly affects the low and high gamma components (p_low gamma_= 0.004, p_high gamma_= 0.021; t-statistic_low gamma_= 2.92 and t-statistic_high gamma_= 2.27). These findings highlight the differential contributions of frequency and probe optimality to neuronal gain modulation across distinct frequency bands in the sample neuron. This single-neuron analysis illustrates the method we will use to examine the impact of visual probe and component frequency on firing rates for the population.

**Figure 3.**
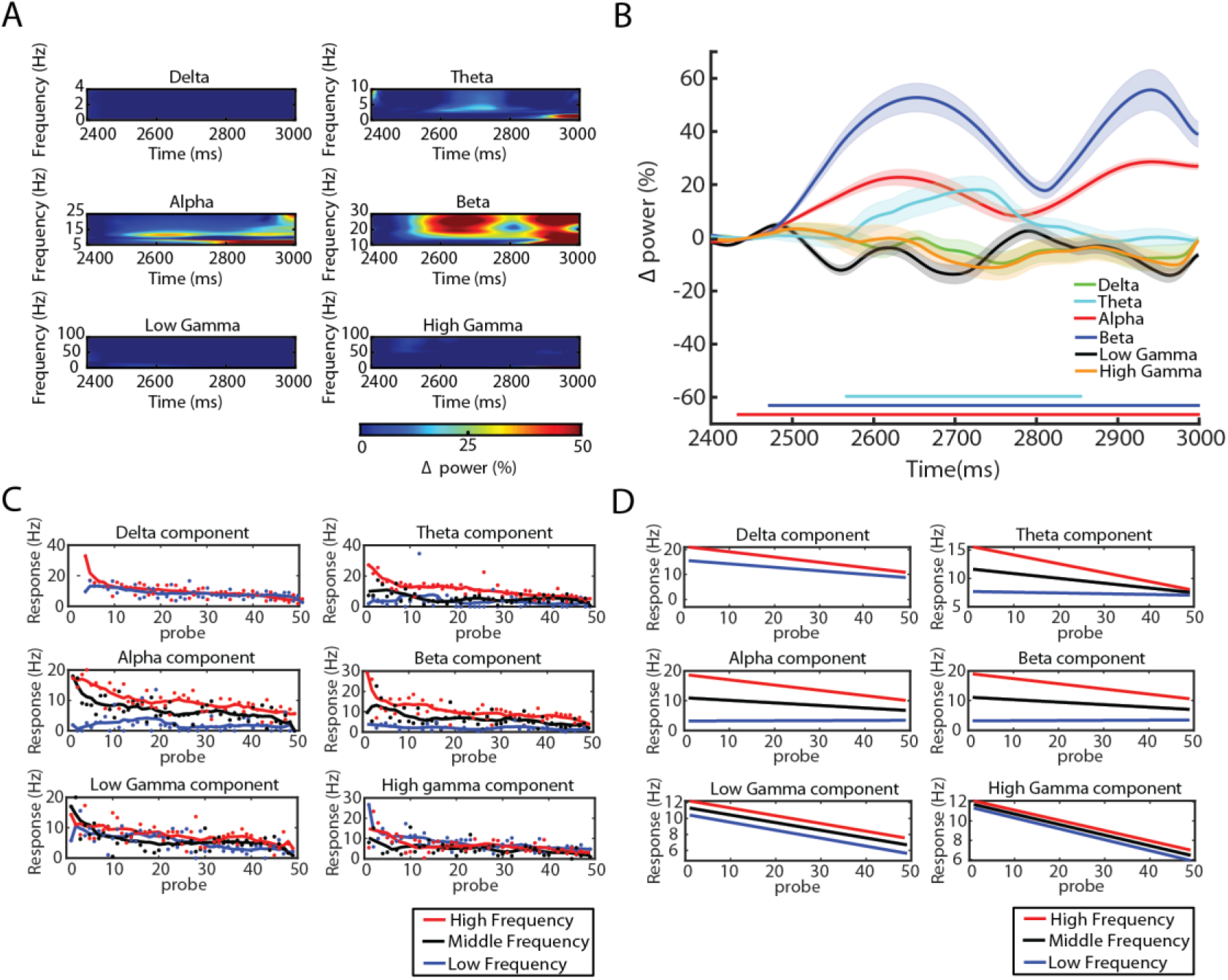
Power of components, frequency-dependent gain modulation, and regression model for the sample neuron. (A) Power differences between IN and OUT conditions during the memory period (2400–3000 ms) for the sample neuron shown in Figure 1B. Each subplot represents one component (delta, theta, alpha, beta, low gamma, and high gamma), illustrating the power of the signal across frequency and time. The plotted values reflect the subtraction of power in the memory IN condition from the memory OUT condition. (B) Averaged power across frequencies for different components during the memory period (2400–3000 ms) for the same sample neuron. The plots depict the power differences of the decomposed components over time, calculated as the power in the memory IN condition subtracted from the memory OUT condition. Solid lines on the bottom indicate significant changes in power during that time bin for the corresponding frequency. In B-D, shading shows the standard error of the mean across trials. (C) Firing rate as a function of ranked probes based on responses during the memory IN condition for the sample recording channel shown in Figure 1B. Each subplot corresponds to one component and illustrates firing rate changes across different frequency groups. Here and in D, frequencies are categorized into high-frequency (red line), middle-frequency (black line), and low-frequency (blue line) groups. (D) Linear regression models fitted to the data presented in Figure 3C. The fitted models illustrate the relationship between firing rate changes and probe optimality for different frequency groups across each decomposed component.

We used the approach demonstrated in Figure 3 to analyze data from 256 recording channels and 131 single neurons. Figure 4A illustrates the power differences for each frequency component during the memory period (2400–3000 ms) between IN and OUT conditions, averaged across all recording channels. This analysis revealed prominent modulations in the power of the alpha, beta and theta components, aligning with the findings from the sample neuron in Figure 3A and demonstrating consistency across the population. Figure 4B provides a cross-sectional view of the population-averaged power differences for each component (see Methods), showing significant changes in the alpha, beta and theta component power during much of the memory period. Figure 4C investigates the influence of frequency on visual signal gain by ranking visual probes from 1 to 49 based on their evoked responses during the memory IN period for 131 neurons. The firing rates associated with these ranked probes were categorized into three frequency groups (high, middle, and low) for each frequency band, based on the frequency of the decomposed components of the corresponding LFP signal on individual trials (see Methods). Across the 6 components, the proportion of neurons showing statistically significant effects (p < 0.05) ranged from 7% to 81%. The alpha, beta, and theta components exhibited the highest proportion of significant neurons (81%, 65%, and 61%, respectively, ANOVA), while the low gamma, high gamma, and delta components showed the lowest proportions (51%, 42%, and 7%, respectively, ANOVA). The mean F-statistics varied across components, with the alpha, theta, and beta components having the highest mean F-statistics (f-statistic_alpha-IN_= 27.51 ± 0.37, f-statistic_theta-IN_= 24.21± 0.54 and f-statistic_beta-IN_= 23.32± 0.21, ANOVA). In contrast, the low gamma, high gamma, and delta components had the lowest mean F-statistics (f-statistic_low gamma-IN_= 6.01 ± 0.54, f-statistic_high gamma-IN_= 3.91± 0.37 and f-statistic_delta-IN_= 2.71 ± 0.10, ANOVA). The population-level analysis reveals that the frequency of the components significantly impact signal gain, with firing rates modulated by frequency in the alpha, beta, and theta components. Figure 4D presents the corresponding analysis for the memory OUT condition, showing how different frequency groups and probe optimality contribute to the modulation of neuronal gain. Across the 6 components, the proportion of neurons showing statistically significant effects of component on frequency (p < 0.05) ranged from 5% to 88%. The alpha, beta, and theta components exhibited the highest proportion of significant neurons (88%, 66%, and 59%, respectively, ANOVA), while the low gamma, high gamma, and delta components showed the lowest proportions (53%, 41%, and 5%, respectively, ANOVA). The mean F-statistics varied across components, with the alpha, theta, and beta components having the highest mean F-statistics (f-statistic_alpha-OUT_= 33.75 ± 2.99, f-statistic_theta-OUT_= 27.91± 2.45 and f-statistic_beta-OUT_= 26.21± 1.48, ANOVA). In contrast, the low gamma, high gamma, and delta components had the lowest mean F-statistics (f-statistic_low gamma-OUT_= 5.70 ± 0.51, f-statistic_high gamma-OUT_= 3.51± 0.34 and f-statistic_delta-OUT_= 2.40 ± 0.13, ANOVA). Notable contributions were observed in the theta, alpha, and beta components. However, in the theta component, the firing rates in the middle- and low-frequency groups were closer together, and a similar trend was observed for the alpha and beta components compared to the memory IN condition. To evaluate how frequency and probe optimality statistically affect the gain of visual signals, a linear regression model was fitted to firing rate changes (as shown in Figure 3D) for each neuron. Figure 4E illustrates the results for the memory IN condition, averaged across 131 neurons. The models reveal the relationship between firing rate changes and probe optimality across different frequency bands (high, middle, and low) and decomposed components (delta, theta, alpha, beta, low gamma, and high gamma). Frequency-specific modulations in firing rates are most significant in the alpha, theta and beta components. Similar trends are observed in the memory OUT condition, as shown in Figure 4F, with alpha, theta and beta bands demonstrating the most significant frequency modulation.

**Figure 4.**
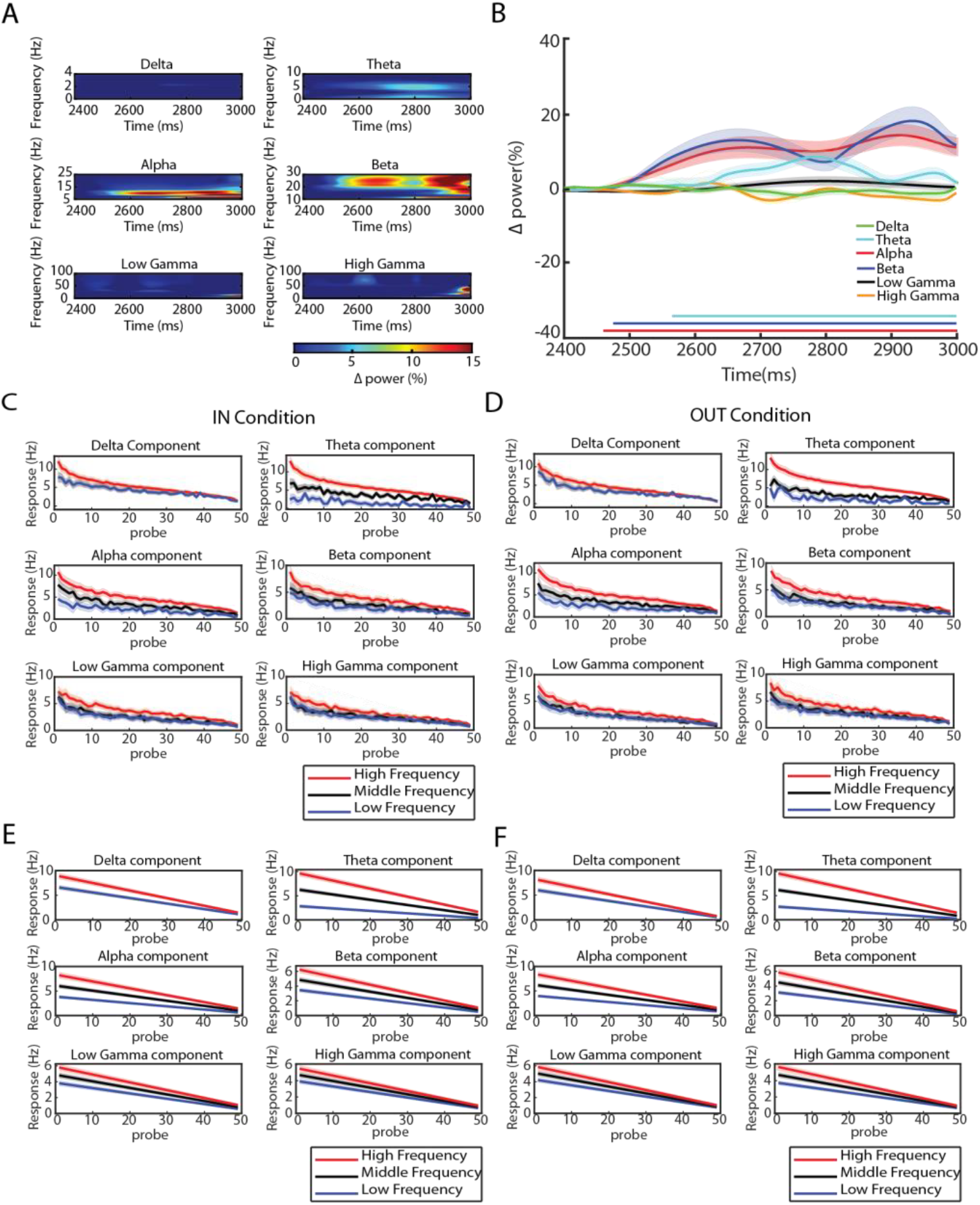
Power of components and frequency-dependent gain modulation across the population. (A) Power differences between IN and OUT conditions during the memory period (2400–3000 ms) across 256 recording channels. Each subplot represents one component (delta, theta, alpha, beta, low gamma, and high gamma), illustrating the power of the signal across frequency and time. The plotted values reflect the subtraction of power in the memory IN condition from the memory OUT condition. (B) Averaged power across frequencies for different components during the memory period (2400–3000 ms) across 256 recording channels. The plots depict the power differences of the decomposed components over time, calculated as the power in the memory IN condition subtracted from the memory OUT condition. Time bins with significant differences from baseline (0-50ms) are indicated by solid lines of the corresponding color below. In B-F, shading shows the standard error of the mean across trials. (C) Averaged firing rate as a function of ranked probes based on responses across 131 neurons during the memory IN condition. Each subplot represents one component and depicts the averaged firing rate changes across different frequency groups. In C-F, frequencies are categorized into high-frequency (red line), middle-frequency (black line), and low-frequency (blue line) groups. (D) Plots same as C, for the memory OUT condition. (E) Averaged linear regression models fitted to the firing rate data of 131 neurons during the memory IN period. The models depict the relationship between firing rate changes and probe optimality for different frequency groups (high, middle, and low) across each decomposed component. (F) Plots same as E, for the memory OUT condition.

To quantify the statistical contributions of frequency, probe optimality, and their interaction, t-statistics were computed from the regression models. Figure 5A displays the averaged t-statistics for the memory IN condition. For the frequency factor (top panel), the strongest modulation occurs in the alpha component (t-statistic_alpha-IN_=5.96±0.43), followed by the theta (t-statistic_theta-IN_= 4.82±0.58) and beta (t-statistic_beta-IN_= 3.24±0.20) components. Probe optimality (middle panel) primarily affects firing rates in the high gamma and low gamma components (t-statistic_high gamma-IN_=1.81±0.11 and t-statistic_low gamma-IN_=1.50±0.12), with a decreasing trend toward the delta component. The interaction between frequency and probe optimality (bottom panel) mirrors the frequency trends, with the alpha component showing the strongest modulation (t-statistic_alpha-IN_=2.89±0.24), followed by theta (t-statistic_theta-IN_=2.30±0.46) and beta (t-statistic_beta-IN_=1.61±0.13). Figure 5B presents the averaged t-statistics for the memory OUT condition, showing a similar pattern to the IN condition. Frequency-related modulation is most significant in the alpha, theta and beta components (t-statistic_alpha-OUT_= 7.68±0.37, t-statistic_theta-OUT_=5.11±0.58, and t-statistic_beta-OUT_=3.50±0.21). Probe optimality (middle panel) predominantly influences the high and low gamma components (t-statistic_high gamma-OUT_=1.87±0.11 and t-statistic_low gamma-OUT_=1.46±0.11). The interaction of frequency and probe optimality (bottom panel) also exhibits a consistent trend, with the alpha, theta and beta components displaying the highest modulation (t-statistic_alpha-OUT_=3.98±0.24, t-statistic_theta-OUT_=2.51±0.33, and t-statistic_beta-OUT_=1.82±0.13). Figures 5C and 5D present histograms showing the number of neurons (out of 131) with significant effects (p < 0.05) of frequency, probe optimality, and their interaction across decomposed components. For instance, in the memory IN condition, more than 76% of neurons (100 out of 131 neurons) exhibit significant frequency modulation in the first four components, aligning with the trends observed in Figures 5A and 5B. This analysis highlights the critical roles of frequency-specific dynamics and probe optimality in modulating visual signal processing across memory conditions. To compare the effects between the memory IN and OUT conditions, Figure 5E shows scatter plots of the slopes from the linear regression models fitted to firing rate changes of 131 neurons for each frequency group across all components. Red dots represent high frequency, black dots represent middle frequency, and blue dots represent low frequency. No significant differences were observed between the IN and OUT conditions across any component (p > 0.05). Therefore, the slight difference in gain between the IN and OUT conditions could result from the higher frequency observed in the IN condition (Figure 2E&F), or a common neural mechanism could drive the variability in both the frequency and the gain.

**Figure 5.**
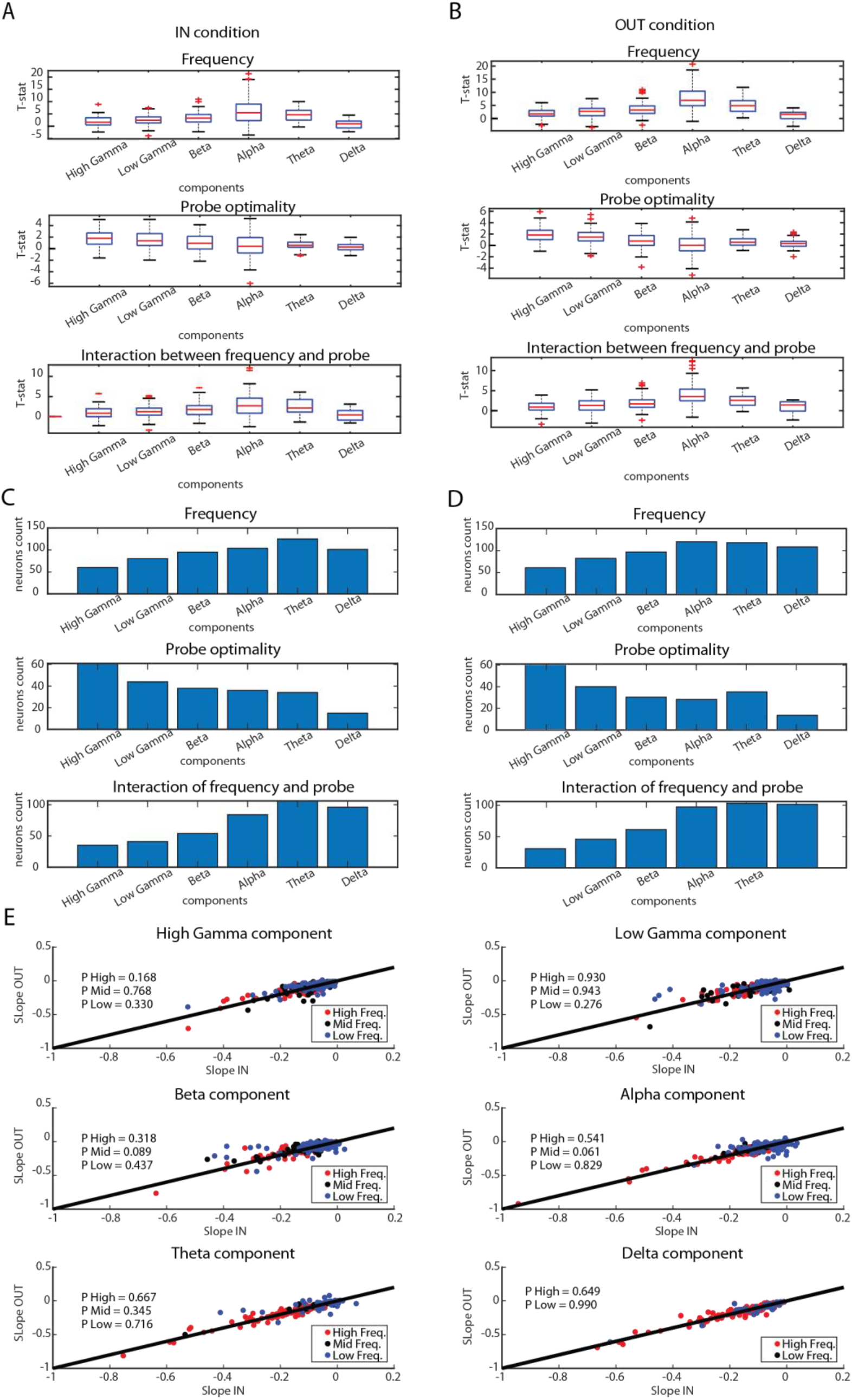
Regression model and statistical analysis. (A) Averaged t-statistics of the linear regression models (n = 131) for the memory IN condition, highlighting the contribution of each factor to firing rate changes across components. These factors include frequency (top panel), probe optimality (middle panel), and the interaction between frequency and probe optimality (bottom panel). The averaged t-statistics illustrate the significance of these factors in modulating firing rates across different frequency bands (delta, theta, alpha, beta, low gamma, and high gamma). (B) Same as A for the memory OUT condition. (C) The number of neurons for which the contributions of each factor—frequency (top panel), probe optimality (middle panel), and the interaction between both factors (bottom panel)—were statistically significant (p < 0.05) in modulating firing rate changes during the memory IN condition. (D) Same as C for the memory OUT condition. (E) Scatter plot showing the slopes of the fitted linear regression models to firing rate changes of 131 neurons for each frequency group and component, for the IN vs. OUT conditions. Red dots represent high frequency, black dots represent middle frequency, and blue dots represent low frequency.

## Discussion

Our study demonstrates that WM selectively modulates both LFP oscillations and spiking activity in the extrastriate cortex, with distinct frequency-dependent effects on sensory processing. Using a memory-guided saccade task in monkeys, we found that WM enhances theta, alpha, and beta power during the memory period while leaving gamma power unchanged. Crucially, after decomposing LFP signals using the MODWT, we revealed that the precise frequency of oscillatory components is correlated with firing rate across trials. Specifically, the frequency of alpha, beta, and theta oscillatory components showed the strongest influence on gain modulation, while gamma frequency was less related to visual gain. Previous work had reported both WM-driven changes in visually-evoked responses, oscillatory power and frequency in extrastriate areas^1–3^. This finding expands on prior results by demonstrating a close link between oscillatory frequency and firing rate on individual trials. These findings suggest that WM-driven oscillatory frequency may determine firing rate, or that variability in frequency and firing rate during WM are produced by a shared neural mechanism.

On a methodological level, the finding that component frequency correlates with gain across trials is reassuring that the MODWT process is isolating meaningful trial-to-trial variation in component properties. Signal decomposition of LFPs has more often focused on spatial decomposition of linear array data to separate modulation based on cortical layers^9,19^. Our use of MODWT builds on recent advances in separating periodic and aperiodic neural components^4^, allowing us to attribute WM effects to specific oscillatory bands while controlling for broadband changes, an important potential confounding factor when interpreting changes in LFPs^21^. Different methods may be best suited to different analyses; the MODWT method as implemented here, for example, doesn’t provide information on variability in component power over time within individual trials, which would be necessary to evaluate ‘bursts’ of oscillatory power^22–24^. This method of identifying components on individual trials may prove useful for increasing the accuracy of WM-related phase coding of information^3^, identifying patterns of functional connectivity between networks of simultaneously recorded areas, and generally understanding with greater precision how cognitive tasks modulate oscillatory signals throughout the brain.

The dissociation between gamma and lower-frequency effects in our study reinforces other recent findings regarding different roles for these frequency ranges. Several lines of evidence suggest that gamma primarily mediates feedforward signaling while alpha/beta rhythms carry feedback information^12,14,15,25–27^. Theta frequencies have been also associated with WM performance in a both human^28–31^ and NHP studies^32,33^; whether theta oscillations play a fundamentally distinct role from alpha/beta frequencies is a question which will need to be addressed in future work. The results presented here are consistent with the broad framework of gamma-feedforward vs. low frequency feedback, with lower-frequency components closely associated with visual gain due to WM-related feedback, while gamma is unmodulated and firing rates primarily reflect probe optimality.

In the absence of firing rate changes, WM induces oscillations within extrastriate areas [Ref Neuron] allowing them to express their representation in the form of a phase code [eLife Mohsen]. The idea that prefrontal WM signals can recruit sensory areas through oscillatory coherence [Prog Neurob] can explain the presence of WM-induced oscillations without drastic firing rate changes in sensory areas; however, it also reveals the challenge that the recorded LFPs from a single area might be the result of its recruitment by several other areas involved in performing a specific task. Therefore, a simple bandpass filtering of LFPs might not be able to reveal the communicated information, in the form of a phase code, between the recorded area and other areas. The current paper is a step toward establishing the significance of LFP decomposition in order to unentangle various sources of oscillation within an area with the ultimate goal of understanding how coherent oscillations between areas can create specific channels of communication and how to read the communicated information at the level of a single trial.

## Methods

### EXPERIMENTAL MODEL DETAILS

#### General and surgical procedures

The data were collected at Montana State University and were used in previous publications ^1,2^. Two adult male rhesus monkeys (Macaca mulatta) were used in this study. All experimental procedures were in accordance with the National Institutes of Health Guide for the Care and Use of Laboratory Animals and the Society for Neuroscience Guidelines and Policies. The protocols for all experimental, surgical, and behavioral procedures were approved by the Montana State University Institutional Animal Care and Use Committee. All surgical procedures were carried out under Isoflurane anesthesia and strict aseptic conditions. Prior to undergoing behavioral training, each animal was implanted with a stainless steel headpost (Gray Matter Research, Bozeman MT), attached to the skull using orthopedic titanium screws and dental acrylic. Following behavioral training, custom-made PEEK recording chambers (interior 22×22 mm) were mounted on the skull and affixed with dental acrylic. Within the chambers two 22×22 mm craniotomies were performed above the prefrontal and extrastriate visual areas (prefrontal chambers were centered at 42 mm A/P, 23 mm M/L and 28 mm A/P, 23 mm M/L; extrastriate craniotomies were centered at −6 mm A/P, 23 mm M/L and −13 mm A/P, 23 mm M/L).

#### Behavioral monitoring

Animals were seated in a custom-made primate chair, with their head restrained and a tube to deliver juice rewards placed in their mouth. Eye position was monitored with an infrared optical eye tracking system (EyeLink 1000 Plus Eye Tracker, SR Research Ltd, Ottawa CA), with a resolution of < 0.01^°^ RMS; eye position was monitored and stored at 2 KHz. The EyeLink PM-910 Illuminator Module and EyeLink 1000 Plus Camera (SR Research Ltd, Ottawa CA) were mounted above the monkey’s head, and captured eye movements via an angled infrared mirror. Juice was delivered via a syringe pump and the Syringe PumpPro software (NE-450 1L-X2, New Era Pump Systems, Inc., Farmingdale NY). Stimulus presentation and juice delivery were controlled using custom software, written in MATLAB using the MonkeyLogic toolbox (Asaad et al., 2013). Visual stimuli were presented on an LED-lit monitor (Asus VG248QE: 24in, resolution 1920×1080, 144 Hz refresh rate), positioned 28.5 cm in front of the animal’s eyes. A photodiode (OSRAM Opto Semiconductors, Sunnyvale CA) was used to record the actual time of stimulus appearance on the monitor, with a continuous signal sampled and stored at 32 KHz.

### Behavioral tasks

#### Eye calibration

The fixation point, a ∼1 dva white circle, appeared in the center of the screen, and the monkey maintained its fixation within a ±1.5 dva window for 1.5 s. For eye calibration, the fixation point could appear either centrally or offset by 10 dva in the vertical or horizontal axis. The monkey was rewarded for maintaining fixation.

#### Preliminary receptive field mapping

Preliminary receptive field mapping was conducted by having the monkey fixate within a ±1.5 dva window around the central fixation point, while ∼2.5×4 dva white bars swept in 8 directions (4 orientations) across the approximate location of the neuron’s receptive field. Responses from the recording site were monitored audibly and visually by the experimenter, and the approximate boundaries of the receptive field were noted for the positioning of stimuli in subsequent behavioral tasks.

#### Memory guided saccade task with visual probes

Monkeys fixated within a ±1.5 dva window around the central fixation point. After 1 s of fixation, a 1.35 dva square target was presented and remained onscreen for 1 s. The animal then remembered the target location while maintaining fixation for 1 s (memory period) before the central fixation point was removed. The animal then had 500 ms to move his eyes to a ± 4 dva window around the previous target location, and remain fixating there for 200 ms to receive a reward. A series of brief (200 ms) visual probes (∼1 dva white circles) appeared in a 7×7 dva grid of locations (1-2.5 dva spacing), both before target presentation (fixation receptive field mapping) and during the memory period (memory period receptive field mapping). Four probes were presented in succession on each trial, with an inter-probe interval of 200 ms. This 7×7 grid of probes was positioned to overlap with the receptive field of the recorded neuron based on the preliminary receptive field mapping described above. The location of the remembered target could vary with respect to the receptive field of recorded neurons (see Fig. 1B).

### Neurophysiological recording

Neurophysiological data was recorded from two adult male rhesus monkeys (Macaca mulatta) across 16 sessions (6 sessions from monkey 1 and 10 sessions from monkey 2). This dataset was previously used in Bahmani et al., 2018.

The electrode was mounted on the recording chamber and positioned within the craniotomy area using a Narishige two-axis platform allowing continuous adjustment of the electrode position. For array electrode recordings a 28-gauge guide tube was lowered as described, and the 16-channel linear array electrode (V-probe, Plexon, Inc., Dallas, TX; distance between electrode contacts was 150mm) was advanced into the brain using the hydraulic microdrive. The array electrode was connected to a headstage pre-amplifier (Neuralynx, Inc., Bozeman MT). Neuralynx Digital Lynx SX and associated software were used for data acquisition. Spike waveforms and continuous data were digitized and stored at 32 kHz for offline spike sorting and data analysis. Spike waveforms were sorted manually, and the quality of isolations for simultaneously recorded neurons confirmed using a support vector machine classifier (see Bahmani et al. 2018 for analysis). Area MT was identified based on stereotaxic location, position relative to nearby sulci, patterns of gray and white matter, and response properties of units encountered. The location of brain areas within the recording chamber was verified via single-electrode exploration prior to beginning data collection with the electrode arrays.

## METHOD DETAILS

### Data Analysis

256 LFP channels were recorded across 16 recording sessions. 131 channels had well-isolated single unit receptive fields. 125 channels were excluded from further analysis due to a lack of strong visual signals.

All analyses were carried out using MATLAB. All population statistics are reported as mean ± SEM (standard error of mean). Statistics are all Wilcoxon signed rank, unless otherwise mentioned.

### Mutual Information

Using the MGS task with probes described above, we calculated the mutual information between probe location and the phase of ongoing alpha-beta oscillations at the time of visually evoked spikes during the memory period. The mutual information between stimulus and response quantifies the ability to discriminate between different stimuli (in this case, different probe locations) based on the response (in this case, based on the alpha-beta phase at the time of spikes). The response window consisted of the 0-200 ms after probe onset, and only spike and LFP data from this period was used in calculations. Next, we used a Hilbert transform to calculate the analytic signal of filtered LFP. Instantaneous phases of each component were quantified by calculating the angles corresponding to the analytic signal. We divided the phase values from –pi to pi into 4 bins of size pi/2 (Panzeri et al., 2007). In order to quantify the mutual information, the first step is measuring the response variability. The most common method for measuring response variability is Shannon entropy (Shannon, 1948). The entropy of the response is

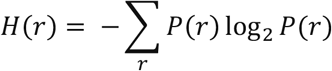

Where *P*(*r*) is the probability of observing response *r* to any stimulus across all trials. The next step is a calculation of the conditional response variability as follows:

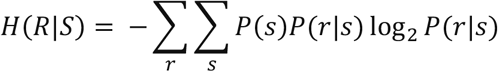

Where *P*(*s*) is the probability of presentation of stimulus s and *P*(*r*|*s*) is the probability of observing response r given presentation of stimulus s. Finally, the mutual information is the difference between response entropy and conditional entropy (Panzeri et al., 2007).

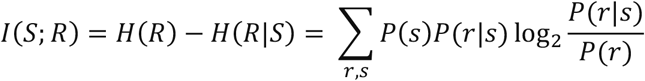

Where *I*(*S*; *R*) is the mutual information between the stimulus space and the response space. Calculations were done either in a space of 49 stimuli (for all 49 probe locations; Figure. 1E). For the shuffling procedure, the spike trains and LFP signals were shuffled across trials and the mutual information is calculated as above

### Spectral Analysis and Peak frequency Detection

To identify oscillator components in single trial LFP signals, we employed the spectral decomposition framework described by Donoghue et. al. (2020). For each trial, the power spectral density (PSD) was estimated using Welch’s averaged periodogram method after notch filter data at 60 Hz to eliminate line noise. The resulting PSD was converted to a decibel scale and smoothed with a Gaussian kernel to attenuate high-frequency noise while preserving the structure of potential oscillatory peak. To isolate narrowband oscillations from the inherent 1/f (“pink noise”) background, an exponential function was fitted to the smoothed PSD. This 1/f fit was subtracted from the spectrum, producing a flattened residual that amplified oscillatory peaks relative to the aperiodic baseline (for further detail see Donoghue et. al. 2020).

Significant peaks were identified as local maxima in the flattened spectrum that exceeded a prominence threshold, a metric quantifying their height relative to neighboring troughs. To avoid spurious detection, peaks were constrained to physiologically relevant frequency bands (e.g., 1-90 Hz). The total number of peaks per trial was recorded, and the mean across all trials was calculated. This average value (6) directly informed the selection of decomposition levels (L) for the subsequent MODWT, ensuring that wavelet scales aligned with the dominant oscillatory components. Parameters such as the smoothing bandwidth (σ) and prominence threshold were validated using synthetic signals with known oscillatory features superimposed on 1/f noise, as described in Donoghue et. al. (2020).

### MODWT Decomposition and Component-Specific Spectral Analysis

To isolate oscillatory dynamics within single-trial LFP signals (0–3200 ms epochs), we applied MODWT using a Frazier-Keiser FK14 basis. The MODWT was chosen for its shift-invariance, tolerance of non-stationary signals, and avoidance of decimation artifacts inherent to classical wavelet transforms. Each trial was decomposed into 5 levels and one approximation component, a parameter selected to align with the average number of oscillatory components identified in the spectral analysis. Wavelet coefficients for each level were reconstructed into time-domain component signals using the inverse MODWT, preserving the temporal resolution of the original data.

For each reconstructed component (levels 1–5), oscillatory properties were quantified by computing the power spectral density (PSD) using Welch’s method with parameters consistent with the initial spectral analysis. The peak frequency of a component was defined as the frequency corresponding to the maximum amplitude within its PSD, restricted to the theoretical frequency band of the decomposition level. For example, a component at level 3 (theoretical band: e.g., 15-30Hz) was analyzed for peaks within a slightly expanded range (e.g., 13–32 Hz) to accommodate spectral leakage. This band-restricted approach ensured that identified peaks aligned with the MODWT’s inherent bandpass structure.

### Power Analysis of Decomposed Components

Following the decomposition process, we analyzed power differences for each frequency component during the memory period (2400–3000 ms) between the IN and OUT conditions. For this analysis, the time-frequency power spectrum of each component (0–3200 ms) was computed for every trial using the continuous wavelet transform (CWT) in MATLAB with a Morlet wavelet. The power spectra were then averaged across trials for each channel. Next, we calculated the percent power change relative to a first 50 ms fixation period as baseline (*[0 to 50 ms]*), separately for the IN and OUT conditions. The condition-specific power changes were then compared by computing the difference (IN minus OUT) for each component.

Additionally, to examine the spectral profile of these power differences, we calculated a cross-sectional view by averaging the power spectra within six predefined frequency bands: delta (1–2 Hz), theta (4–6 Hz), alpha (8–12 Hz), beta (18–24 Hz), low gamma (36–44 Hz), and high gamma (76–84 Hz). This allowed us to compare condition-specific power modulation across distinct oscillatory ranges.

### Gain Modulation Analysis by Frequency Band

After decomposing the LFP signals into six oscillatory components, we examined how frequency-specific activity modulates visual signal gain. For this analysis, we segmented both the decomposed components and their corresponding spike trains into 230-ms epochs (from 30 ms before to 200 ms after probe onset). This time window captured the neural dynamics surrounding visual probe presentation while minimizing contamination from preceding events.

We quantified frequency-dependent gain modulation through a multi-step process. First, we ranked all 49 visual probes for each neuron based on their evoked firing rates during the memory IN period (0 to 200 ms). This ranking established a continuum of probe effectiveness in driving neural responses. Next, we categorized these ranked probes according to the concurrent power of each LFP frequency component, creating distinct power groups for analysis.

For theta (4-6 Hz), alpha (8-12 Hz), beta (18-24 Hz), low gamma (36-44 Hz), and high gamma (76-84 Hz) bands, we established three power groups: high-power trials (exceeding the 66th percentile of the power distribution), medium-power trials (between the 33rd and 66th percentiles), and low-power trials (below the 33rd percentile). Due to the characteristically slower dynamics of delta activity (1-2 Hz), we used a two-group division with high-power trials (above the 50th percentile) and low-power trials (below the 50th percentile).

This grouping approach allowed us to construct separate firing rate curves for each frequency (power) condition while maintaining the probe optimality ranking. By comparing these curves across frequency (power) groups within each frequency band, we could assess how different levels of oscillatory activity modulated the gain of visual responses. The analysis preserved the trial-by-trial relationship between LFP power and spiking activity while controlling for probe effectiveness.

All statistical comparisons were performed using ANOVA with appropriate corrections for multiple comparisons across frequency bands. The analysis was implemented in Matlab (MathWorks) using custom scripts that are publicly available [repository information].

### Regression Analysis of Frequency-Specific Modulation

To systematically examine how oscillatory frequency components and probe optimality influence neural responses, we conducted linear regression analyses on our frequency-divided dataset. For each frequency band (delta: 1-2 Hz, theta: 4-6 Hz, alpha: 8-12 Hz, beta: 18-24 Hz, low gamma: 36-44 Hz, and high gamma: 76-84 Hz), we implemented separate regression models using MATLAB’s *fitlm* function for each neuron. The models characterized how both the frequency band power and probe optimality collectively shaped firing rate patterns.

The regression framework for each frequency band took the form:

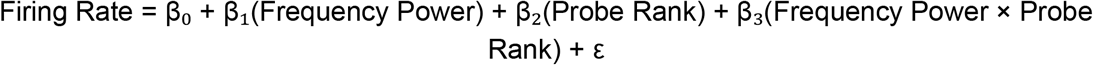

We carefully validated model assumptions by examining residual distributions and conducting diagnostic tests. This included verifying linearity through partial regression plots, assessing normality of residuals using Kolmogorov-Smirnov tests, and checking for heteroscedasticity with White’s test. Any trials exhibiting excessive leverage (Cook’s distance > 3× the mean) were excluded from the final analysis to ensure robust parameter estimates.

The regression coefficients revealed distinct patterns of frequency-specific modulation:

‐ Significant β_1_ coefficients indicated that firing rates systematically varied with the strength of particular frequency components
‐ β_2_ coefficients quantified the expected probe rank selectivity
‐ Significant interaction terms (β_3_) demonstrated how frequency power modulated the relationship between probe optimality and neural response

All statistical tests employed false discovery rate correction (q < 0.05) to account for multiple comparisons across frequency bands. The complete analysis pipeline, implemented in MATLAB 2022b, has been made publicly available to ensure reproducibility. This rigorous approach provided quantitative insights into how different frequency ranges participate in gain modulation of visual responses.

## QUANTIFICATION AND STATISTICAL ANALYSIS

Statistical analysis was performed using custom code written in Matlab (MathWorks). Statistical details including means, p-values, exact value of n, and what n represents are described in the Results; p and n values for plotted data are also shown on the relevant figure panel. All population statistics are reported as mean ± SE (standard error). Nonparametric statistical tests (Wilcoxon signed-rank for paired comparisons, Wilcoxon rank sum for unpaired comparisons) are used throughout for calculating p values.

